# Connexin43 Overexpression Promotes Bone Regeneration by Osteogenesis and Angiogenesis in Rat Glucocorticoid-Induced Osteonecrosis of the Femoral Head

**DOI:** 10.1101/2022.06.13.496037

**Authors:** Xin Zhao, Changjun Chen, Yue Luo, Donghai Li, Qiuru Wang, Yuying Fang, Pengde Kang

## Abstract

Glucocorticoids induced osteonecrosis of the femoral head (GIONFH) is a devastating orthopedic disease. Previous studies suggested that connexin43 (Cx43) is involved in the process of osteogenesis and angiogenesis. However, the role of Cx43 potentiates in the osteogenesis and angiogenesis of bone marrow-derived stromal stem cells (BMSCs) in GIONFH is still not investigated. In this study, BMSCs were isolated and transfected with green fluorescent protein (GFP) or the fusion gene encoding GFP and Cx43. The osteogenic differentiation of BMSCs were detected after transfected with Cx43. In addition, the migration abilities and angiogenesis of human umbilical vein endothelial cells (HUVECs) were been detected after induced by transfected BMSCs supernatants in vitro. Our results showed that Cx43 overexpression in BMSCs promoted osteogenic differentiation and angiogenesis in vitro. Finally, we established GC-ONFH rat model, then, a certain amount of transfected or controlled BMSCs were injected into the tibia of the rats. Immunohistological staining and micro-CT scanning results showed that the transplanted experiment group had significantly promoted more bone regeneration, vessel volume and the expressions of Runx2, ALP, COL I, VEGF and CD31 when compared with the effects of the negative or control groups. This study demonstrated for the first time that the Cx43 overexpression in BMSCs could promote bone regeneration as seen in the osteogenesis and angiogenesis process, suggesting that Cx43 may serve as a therapeutic gene target for GIONFH treatment.

## Background

Glucocorticoids are important therapeutic agents that have been widely used as anti-inflammatory and immunosuppressive drugs(6, 33, 45, 47). However, their therapeutic benefits are often associated with serious complications, such as osteocyte apoptosis and osteonecrosis(8, 15, 43). Osteonecrosis of the femoral head (ONFH) is a destructive disease that is characterized by cell death within the femoral head, progressive degeneration of the hip joint, and severely lowered quality of life(27, 55). Identified risk factors for ONFH include glucocorticoids use, alcohol consumption and trauma, but its pathogenesis remains poorly understood. Surgical intervention is currently a traditional treatment strategy for ONFH; however, it is an invasive procedure and could influence the patients’ quality of life. Therefore, investigating non-surgical treatment methods for ONFH is necessary.

Bone marrow-derived stromal stem cells (BMSCs) have the potential for self-renewal and multi-directional differentiation and have been widely used in tissue regeneration or repair(4–5, 9, 51, 59). BMSCs are suitable for clinical applications and are easily obtained from patients, and immunological incompatibilities could be avoided with autologous transplantation(18, 30). Preclinical studies showed that bone healing began two weeks after autologous bone marrow stem cell transplantation for ONFH treatment and achieved complete healing after nine weeks(42). A follow-up study of five years showed that the clinical effects of autologous bone marrow cells transplantation combined with autogenous iliac cancellous bone grafts for the treatment of moderate lesion of ONFH are comparable to femoral head-preserving surgeries, suggesting that BMSCs transplantation is a promising method for ONFH treatment(24). Numerous studies in the literature had demonstrated the effectiveness of stem cell transplantation in the treatment of early stages of osteonecrosis(12, 19–20, 57). Hernigou P et al. found that autologous bone marrow transplantation significantly improved the natural course of the early stage of ONFH when compared with simple nucleus pulposus decompression surgery(19). Furthermore, it was found that P-glycoprotein (P-gp)-overexpression BMSC transplantation could decrease the risk of glucocorticoids-induced ONFH (GIONFH)(16), and SDF-1α overexpression of BMSCs could promote osteogenesis and vascularization, therefore reduce the incidence of GIONFH(54); In addition, the elevated expression of BMP-2 and BFGF in BMSCs could accelerate bone repair of ONFH(39). All these findings indicate that there are a variety of factors that play important roles in the therapeutic effects of BMSCs on ONFH, and we believe that transplantation of genetically modified BMSCs can be used as an effective method for the treatment of earlystage GIONFH. Nevertheless, the pathogenesis of GIONFH is not fully understood, and it’s important to further study the precise mechanisms of GIONFH and find new methods to inhibit or delay osteonecrosis occurrence.

Gap junction channels are formed by two hemichannels, which are composed of six transmembrane proteins called connexins(23, 25). It has been reported that these connexins play a vital role in tissue homeostasis(7), in the regulation of cell proliferation and growth, and in cell differentiation and development(1, 40). There are at least 21 different human connexins that have been reported in the literature so far, and they have a series of homologs that showed different tissue or cells specificities(2, 29). Among these connexins, connexin43 (Cx43) is considered to be the main component of gap junctions in hematopoietic tissue(29). Moorer & Stains reported that Cx43 is greatly associated with the process of osteogenesis and osteoblast function(35). Li et al. also reported that Cx43 had the function of regulating extracellular signal-regulated kinase (ERK) activity, therefore, it consequently regulates Runx2, which is an essential transcription factor for osteoblast differentiation(31). In addition, several studies have demonstrated that Cx43 is greatly involved in the process of angiogenesis(49), which is essential for osteogenesis. More importantly, the knockdown of Cx43 expression in the endothelial progenitor cells could decrease the expression of VEGF, and weaken the angiogenic potential of the cells(50). Furthermore, the use of high doses of dexamethasone could inhibit the osteogenesis and angiogenesis in bone tissue(53, 56), and could also down-regulate the expression of Cx43(44). Based on the facts that osteogenesis and angiogenesis are essential for bone regeneration, we speculated that Cx43 may play a critical role in GIONFH treatment. This present work aimed to examine the function of Cx43 in BMSCs-induced osteogenesis and angiogenesis for GIONFH treatment.

## MATERIALS AND METHODS

### Rat BMSCs isolation and transfection

BMSCs were isolated from SD rats (males, four weeks old, body weight 100±15 g) as previously described(63). A total of 1-2 ml bone marrow was aspirated by a heparinized syringe from the lateral tibial tubercle of the rats. The cells were cultured in low-glucose DMEM supplemented with 10% fetal bovine serum (FBS, Gibco, USA) and 1% penicillin–streptomycin in humidified atmosphere of 5% CO_2_ at 37°C. The cells were passaged when they reached 100% confluence, and after three to five passage, the cells were analyzed by flow cytometry and kept for further use.

Third-passage BMSCs were transfected with a lentiviral plasmid carrying the green fluorescent protein (GFP) and Cx43 (GeneChem, Shanghai, China) or a lentiviral plasmid carrying GFP and a negative control sequence (GeneChem, Shanghai, China). The two groups of the transfected cells were called Cx43-GFP-BMSCs and GFP-BMSCs, whereas the non-transfected cells were labeled the Control group. After 12 h of transfection, the medium was replaced with fresh complete medium, and the transfected cells with a density of more than 80% confluency were purified using complete medium containing 3 μg/ml puromycin for 5-6 days.

### Flow cytometry

For phenotypic characterization analysis, 5×10^5^ BMSCs were incubated with fluorescein CD34, CD45, CD29, and CD90 at a dilution rate of 1:100 for 30 min at 4°C, and then flow cytometry analysis was carried out using a flow cytometer (BD FACSAria, USA). FlowJo 7.6.5 software (Tree Star Inc., Ashland, OR, USA) was used for data analysis. The cells that treated by puromycin were suspended with culture medium and subjected for further use.

### Real-time polymerase chain reaction (RT–PCR)

Total RNA was extracted from the BMSCs, HUVECs and the rats’ femoral heads using the TRIzol (Invitrogen, USA) method according to the manufacturer’s instructions. RNA was reverse-transcribed to cDNA according to the manufacturer’s instructions (Thermo, USA). The expression levels of Cx43, Runx2, ALP, and COL I at the mRNA level were measured with IQ^TM^ SYBR Green Supermix (Bio–RAD). The following primers were provided by TsingKe (Beijing, China): Cx43 forward: 5’-CTCACCTTTGTGCCTTCC-3’, reverse: 5’-CTCACCTCCCTGATGCTAA-3’; Runx2 forward: 5’TCGGAAAGGGACGAGAG-3’; reverse: 5’-TTCAAACGCATACCTGCAT-3’; ALP forward: 5’-CCGCAGGATGTGAACTACT-3’; reverse: 5’-GGTACTGACGGAAGAAGGG-3’; COL I: 5’-TGCAAGAACAGCGTAGCC-3’; reverse: 5’-CAGCCATCCACAAGCGT-3’; VEGF forward: 5’-ACAGGGAAGACAATGGGA-3’; reverse: 5’-CTGGAAGTGAGCCAACG-3’; CD31 forward: 5’-TCCCCACCCAAAGTAGC-3’; reverse: 5’-TAAACAGCGCCTCCCAT-3’. PCR was performed for 40 cycles, and the expression levels of mRNAs were calculated by the 2^−ΔΔCt^ method, and GAPDH was used as a reference gene. All experiments were repeatedly performed in three times.

### Western blot analysis

Proteins were extracted using ice-cold RIPA buffer (Beyotime Biotechnology, China) containing 1 mmol/L PMSF. Lysates mixture was then centrifuged at 14000 rpm for 15 min, and the concentration of protein in the supernatant was measured by a BCA Protein Assay kit (Beyotime Biotechnology, China). The proteins samples (20 μg/lane) were separated by SDS–PAGE and transferred onto PVDF membranes (0.45μm, Millipore, USA) using a transfer unit (Bio–RAD, USA). Thereafter, the membranes were blocked in TBS solution containing 5% non-fat milk for 2 h at room temperature, and then incubated with primary antibodies including Cx43 (mouse, 1:1000, abcam), Runx2 (rabbit, 1:1000, Boster Bio), ALP (rabbit, 1:500, Boster Bio), COL I (rabbit, 1:1000, Boster Bio), VEGF (mouse, 1:1000, abcam), CD31 (mouse, 1:1000, abcam) and GAPDH (mouse, 1:1000, Sigma–Aldrich) for overnight at a temperature of 4°C, respectovely. After been washed with TBST, the membranes were incubated with secondary antibody (1:10000, ZSBIO, Beijing, China) for two hours. After been washed with TBST three times, the membranes were visualized by enhanced chemiluminescence (ECL, Beyotime). The digitized images were analyzed using IPP software 6.0 (Media Cybernetic, USA).

### Immunofluorescence staining

The osteoblasts were cultured on small coverslips, and allowed to adhere, and then were treated either with or without 10^−6^ mol/L MPS for four days. Thereafter, the cells in the two groups were washed with PBS solution for 5 min each, then fixed with 4% paraformaldehyde for 30 min, and washed with PBS for 5 min each. The cells were also permeabilized with 0.3% Triton X-100 for 20 min, and washed in PBS for 5 min. Then the cells were blocked for 30 min with 10% goat serum at 37°C, and the cells of both groups were incubated with Cx43 (mouse, 1:200; Santa Cruz, CA, USA), Runx2 (rabbit, 1:300; Boster Bio, China), ALP or COL I (rabbit, 1:1000, Boster Bio) for overnight at 4°C. The next day, the cells were washed in PBS three times, and the cells of both groups were incubated with secondary antibodies (FITC-conjugated anti-mouse, 1:300; AlexaFluor 488-conjugated anti-rabbit, 1:300) for 2 h at room temperature. Finally, samples of both groups were incubated with DAPI for 10 min at room temperature. The images were acquired with a fluorescence microscope (Zeiss, Carl Zeiss, Germany).

### Inducing Osteogenic differentiation

Total of 2×10^4^ cells/cm^2^ were seeded in a 24-well plate precoated with Gelatin solution and allowed to adhere. The cells were then treated with osteogenic differentiation induction medium containing Glutamine, Ascorbate, β-sodium glycerophosphate, and penicillin–streptomycin (Cyagen, Shanghai, China). The induction medium was being replaced every three days. Alizarin red staining assay kit was applied used to detect the mineralization of BMSCs.

### Adipogenic differentiation induction

For achieving adipogenesis, the cells were incubated with the adipogenesis-inducing medium A (Cyagen, Shanghai, China) for three days, and then this medium was replaced with the adipogenesis-inducing medium B (Cyagen, Shanghai, China), continuing incubation which was incubated for two days, and then replaced back with medium A for three days. After 14 days, the cells were harvested and stained with Oil Red O for 30 min.

### Chondrogenic differentiation induction

A total of 6×10^5^ cells were put into a 15-ml sterile centrifuge tube and centrifuged at 250 g for 4 min. The suspension was discarded, and a 0.5ml of chondrogenic differentiation induction medium without TGF-β3 was added to resuspend the cells. Then, the cells were washed by repeating the process above. Thereafter, the cells were treated with chondrogenic differentiation induction medium containing Dexamethasone, Ascorbate, ITS supplement, Sodium pyruvate, Proline, and TGF-β3 for three weeks according to the manufacturer’s instruction (Cyagen, Shanghai, China). Frozen slices of the chondrocytes balls were made and measured by Saffron O staining and observed under a light microscope (Leica, Japan).

### Cell counting kit-8

Cell Counting Kit-8 (CCK-8) assays (Dojindo, Jepan) were used to investigate the cell proliferation. In a brief, the intervened or control cells were trypsinized and replanted into 96-well plates with 5×10^3^ cells/well for the proliferation assay. Then, a 100 μl of DMEM containing 10 μl of CCK-8 working solution was added into each well at the 1, 2, and 3 d time points and incubated for three hours in an incubator. After the incubation, the optical density at 450 nm of absorbance was detected with a microplate reader.

### Cell apoptosis

After overexpression of Cx43 in BMSCs, the cells were collected and incubated with Annexin V-FITC/PI apoptosis detection working solution (Becton Dickinson) according to the instructions (Becton Dickinson). Then, flow cytometry analysis was carried out using Flow cytometer (BD FACSAria, USA) and the FlowJo 7.6.5 software (Tree Star Inc.) was used for data analysis.

### ALP staining

All groups of BMSCs were cultured in 24-well plates at a density of 2×10^4^ cells/well. Ten days after the osteogenic differentiation induction process on different groups of BMSCs, including the control group, GFP-BMSCs and Cx43-GFP-BMSCs, the cells were incubated with BCIP/NBT ALP Color Development Substrate (Beyotime, Shanghai, China) for 20 min. Thereafter, the cells were lysed in radioimmunoprecipitation assay lysis buffer (Sigma–Aldrich) to measure the activity of ALP. ALP activity was evaluated using an ALP assay kit (Nanjing Jiancheng Biotechnology Co., Ltd., Nanjing China). The optical density was measured with a microplate reader at 520 nm.

### Alizarin red staining

After twenty-one days, the osteogenic differentiation induction mediums of all groups were discarded, and gently washed twice with PBS solution. Then, they were fixed with 4% paraformaldehyde for 30 min; staining with alizarin red for 20 min; and finally observed under a light microscope (Olympus, Japan). To quantify the mineralization, the calcium deposits were desorbed using 10% cetylpyridinium chloride (Sigma), and the absorbance at 570 nm was measured.

### Cell migration assay

Cell migration was evaluated by scratch wound assay, which was described in detail in our previous study(21). Briefly, the HUVECs were planted into a 6-well plate at a density of 5×10^5^ cells/well and cultured in growth medium until the confluence of the cells reached 100%, the cells were then scratched with sterile 200μl pipet tips, and the cells debris were removed by PBS solution. Thereafter, the cells were put back into the incubator (37°C, 5% CO_2_) after adding the conditioned medium that containing 2% FBS. The migration of cells was photographed under a microscope with Zen Imaging software after 24 hours of scratching. The method for calculating the percentage of wound healing was in consistent with our previous study(60).

### Tube formation assay

To investigate the effect of Cx43-GFP-BMSCs on the angiogenic differentiation of HUVECs, we collected the culture supernatants from the different treated groups of BMSCs to be kept as conditioned mediums for further uses. HUVECs were seeded into 6-well plates (1×10^5^ cells per well) and cultured with BMSCs-conditioned medium for 3 days. Thereafter, the angiogenic-associated genes, including VEGF and CD31, were detected by Western blot analysis.

To investigate the angiogenic effects of BMSCs-conditioned medium on HUVECs, the tube formation assay was performed. In a brief, Matrigel (50μL/well, Corning, USA) was placed into a 96-well plate on ice and kept for 30 min at 37°C until it became solidified. Subsequently, 2×10^4^ HUVECs/well were seeded on the surface of Matrigel and incubated with either BMSCs-conditioned medium or normal culture medium. Twelve hours later, tube formation was observed with a light microscope (Olympus, Japan). The ability to form capillary-like structures was determined by the number of branch points and tubule lengths in three randomly selected fields under the microscope.

### Animal model and grouping

All animal experiments were performed based on the guidelines that formulated by the National Institution of Health on the humane use and care of laboratory animals, and all animal protocols were granted by the Institutes Animal Care and Use Committee of West China Medical School of Sichuan University. A total of 24 adult male Sprague–Dawley (SD) rats (450-500g) were purchased from the experimental animal center of Sichuan University. All rats were housed at the Animal Center of Sichuan University and kept for eight weeks.

The rats were weighted and randomly divided into four groups, including the normal group (control, non-transplantation), MPS group (GIONFH, non-transplantation), negative control group (transplantation of GFP-BMSCs), and experiment group (transplantation of Cx43-GFP-BMSCs). Based on our previous study(63), we adopted LPS & MPS to establish the ONFH model. Two weeks after the first MPS injection, amount of BMSCs (1×10^7^ GFP-BMSCs or Cx43-GFP-BMSCs) were injected into the tibia of the rats in the intervened groups, whereas the rats in the control or MPS groups had not received any treatment.

### Micro-CT scanning

As described in our previous study, the femoral head morphologic changes were detected by a micro-computed tomography (micro-CT) system (Inveon Multimodality Gantry STD CT) at a resolution of 9 μm with the following parameters: current, 112 μA; x-ray energy, 80 kvp; and exposure time, 370 milliseconds. To evaluate the bone morphological changes in the femoral head, the parameters of new bone volume/total volume (BV/TV), trabecular thickness (Tb. Th), trabecular number (Tb. N), and trabecular separation (Tb,Sp) were calculated. To evaluate the angiogenesis, 3D reconstructions images of blood vessels were obtained, and morphometric parameters, including blood vessel areas and total blood vessel length in the femoral head, were calculated.

### Angiography

Angiography was performed as previously described(48). In a brief, eight weeks after the operation, the rats were anesthetized and perfused with Microfil (Microfil MV-122; Flow Tech, Carver, MA, USA). The method was as followsed: firstly, the hair of the chest was shaved, and the rib cage was opened with a pair of scissors. Then, a 100ml of heparinized saline and 20ml of Microfil were continuously injected into the cardiac apex at a rate of 2ml/min. Finally, the perfused rats were laid flat at 4°C in the refrigerator overnight to ensure complete polymerization. The bilateral femoral heads were dissected and harvested, and decalcified in 10% EDTA solution for about 4 weeks; and the samples were scanned by micro-CT as described above, and the blood vessels in the femoral head were reconstructed using CTVol software.

### Histology and Immunohistochemistry (IHC)

Hematoxylin and eosin (H&E) staining was applied to assess the histomorphological changes in the femoral head. After the rats were sacrificed, their femoral heads were harvested, decalcified, and sectioned in the coronal plane. Some of the sections were subjected to H&E staining to evaluate the trabecular structures. Briefly, after fixed with 10% formalin for 24 h, the femoral heads were decalcified in Ethylene diamine tetraacetic acid (EDTA, 10%) solution for approximate 4 weeks, and then embedded in paraffin. Samples were cut into 4μm thick sections, deparaffinized in xylene, dehydrated in a gradient of ethyl alcohol, and washed 3 times with distilled water. H&E staining was performed to observe the rate of empty osteocytes lacunae and the destruction degree of bone trabecula. The software of Image-Pro Plus 6.0 (Media Cybernetics, Baltimore, MD, USA) was used to count the rate of the empty osteocytes lacunae, and the specific methods were described previously(61).

The expressions levels of Runx2, ALP, COL I, VEGF and CD31 were detected by immunohistochemistry. In a brief, the tissues were embedded with paraffin and conventionally sliced into 4μm thick sections. The sections were baked for 2 h at 60°C before dewaxed in xylene, and rehydration through graded ethanol. Subsequently, they were placed into sodium citrate buffer pH 6.0 and heated up to 100°C for 5 min for antigen repair. After cooling down, the sections were incubated with 5% goat serum (Solarbio, China) at 37°C for 30 min, then the sections were incubated with rabbit anti-rat Runx2 monoclonal antibodies (rabbit, 1:1000, Boster Bio), ALP (rabbit, 1:1000, Boster Bio), COL I (rabbit, 1:1000, Boster Bio), VEGF (mouse, 1:1000, abcam), and CD31 (mouse, 1:1000, abcam) at 4°C for overnight, and lastly incubated with the secondary antibody (1:1000; biotinylated goat anti-mouse lgG, ZSBIO, China) at 37°C for one hour. The sections were developed with diaminobenzidine (DAB; Beyotime Biotechnology, China) to detect the targeted antibody. After staining, the sections were sealed up with balsam before being observed under an optical microscope (Olympus Optical, China). The brownish-yellow that showed color in the cytoplasm or cytomembrane indicated positive results, and other findings indicated negative results. Five fields were randomly selected for detection to calculate the positive expression. The experiments were repeatedly performed three times.

### Statistical analysis

All data were analyzed using SPSS 22.0 software (SPSS, IBM Corporation, USA), and aere presented as the mean ± SD. Statistical significant differences between two groups were analyzed with the Student’s t-test, and the significance among multiple groups was analyzed using one-way ANOVA with Tukey’s post hoc multiple comparison tests, respectively. A correlation analysis was carried out using a two-tailed Spearman’s rank correlation coefficient (r). A P value<0.05 was regarded as statistically significant. All experiments were repeated at least three times.

## Results

### Characterization of BMSCs and transfection efficiency

The characteristics of the isolated cells were determined by flow cytometry method. The cells were positive for CD29 and CD90, but negative for CD34 and CD45 (Fig. 1A), which were used as typical biomarkers for BMSCs. The multi-lineage differentiation potential of BMSCs was detected, as shown in Fig. 1B, BMSCs could be differentiated to osteoblasts, adipocytes and chondrocytes, which confirmed the stemness of BMSCs. Then, BMSCs were infected with lentiviral vectors carrying either the Cx43 gene combined with GFP or only GFP, and the stable transgenic Cx43-GFP-BMSCs and GFP-BMSCs cells were purified using puromycin for 5-6 days, and the results were confirmed by fluorescence (Fig. 1C). The expression of Cx43 was detected at both the mRNA and protein levels, which suggested that the cells were successfully transfected with the Cx43 gene (Fig. 1D-F).

**Figure 1.**
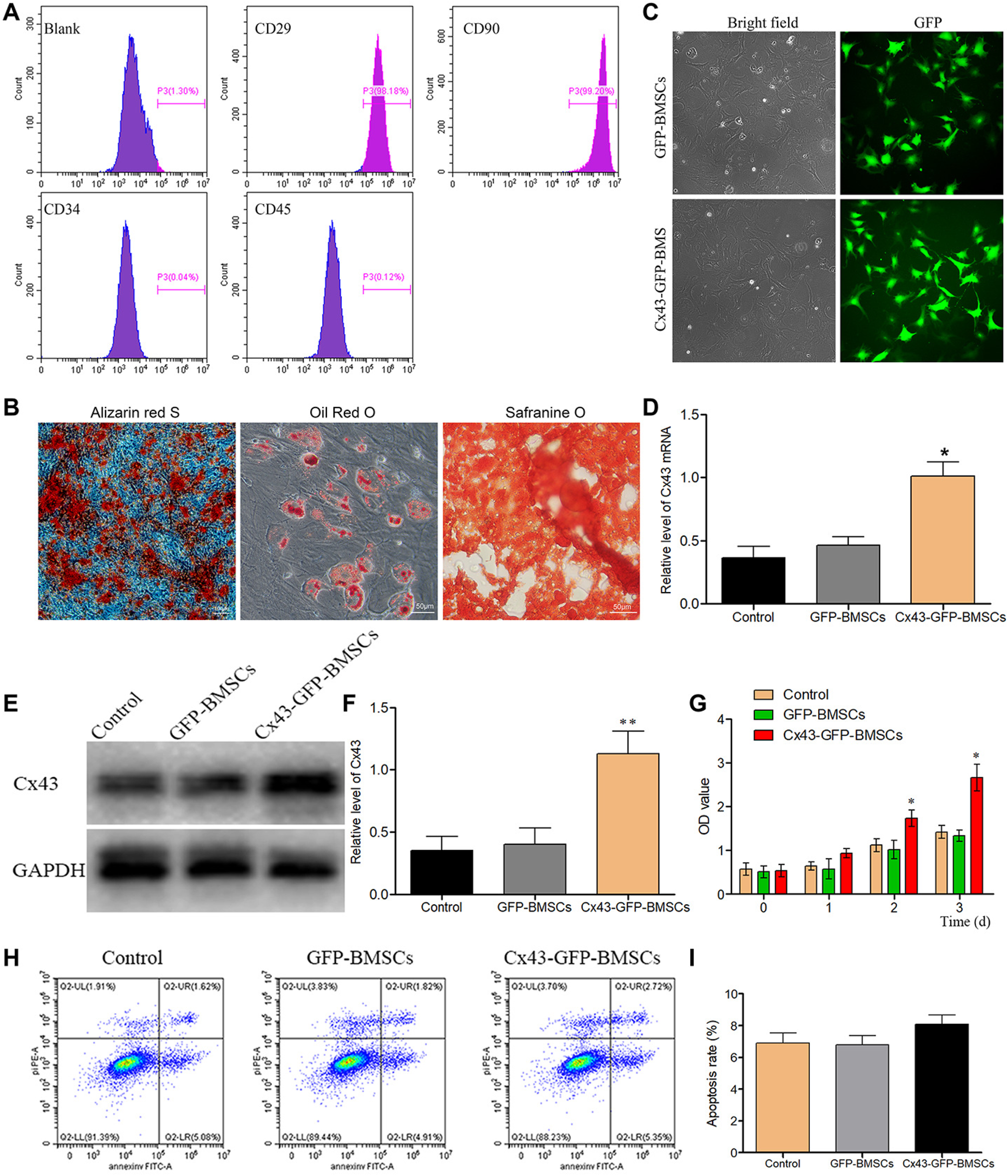
Characteristics of isolated rats’ bone marrow stromal stem cells (BMSCs) and transfection efficacy evaluation. (A) The isolated BMSCs were positive for CD29 and CD90, but negative for CD45 and CD34. The X axis represents fluorescence intensity; (B) BMSCs were differentiated to osteoblasts, adipocytes and chondrocytes; (C) BMSCs were successfully transfected with lentiviral vectors, as indicated by fluorescence; (D) The expression of Cx43 mRNA was detected by RT–PCR; (E) After transfected with the Cx43 gene, the protein expression level of Cx43 was detected by Western blot; (F) Statistical analysis for Cx43 expression; (G) The effects of Lv-Cx43 on BMSCs proliferation. (H) The effects of Lv-Cx43 on BMSCs apoptosis measured by flow cytometry, Q2-UL represents necrotic cells, Q2-UR represents late apoptotic cells, Q2-LR represents early apoptotic cells and Q2-LL represents live cells; (I) Statistical analysis of apoptosis. Each experiment was repeatedly performed at least three times. *P<0.05.

Furthermore, Cx43 was greatly involved in cells proliferation and apoptosis. Therefore, we investigated these processes after Cx43 transfection. As depicted in Fig. 1G, Cx43 transfection in BMSCs had better results on reducing the inhibition effects of MPS on cells proliferation when compared with GFP-BMSCs or non-transfected cells. In addition, the apoptosis rate of cells was not significantly increased in the Cx43-GFP-BMSCs group compared with the GFP-BMSCs or non-transfection group under MPS treatment (Fig. 1H, 1I).

### Cx43 overexpression in BMSCs promotes angiogenesis and endothelial cells recruitment in vitro

Previous studies reported that Cx43 is greatly involved in the process of angiogenesis and endothelial cell recruitment. Therefore, Human umbilical cord vein endothelial cells (HUVECs) were used in this research to evaluate the tube formation activity of transgenic BMSCs. We found that the overexpression of Cx43 in BMSCs had resulted in a significantly higher tube formation ability than GFP-BMSCs or control groups (Fig. 2A-2C). After MPS treatment, which was used to mimic osteonecrosis in vitro, the tube formation ability was decreased (Fig. 2A-2C); However, Cx43 overexpression had significantly improved the tube formation ability compared with MPS treatment alone (Fig. 2A-2C). In addition, the expression of CD31, which is one of the markers of angiogenesis, was detected, and HUVECs were cultured in the supernatants of Cx43-GFP-BMSCs, GFP-BMSCs or non-transfected cells for 4 days. Results demonstrated that the expression of CD31 was significantly increased under the treatment of the Cx43-GFP-BMSCs supernatants in both the absence or presence conditions of MPS (Fig.2D, 2E).

**Figure 2.**
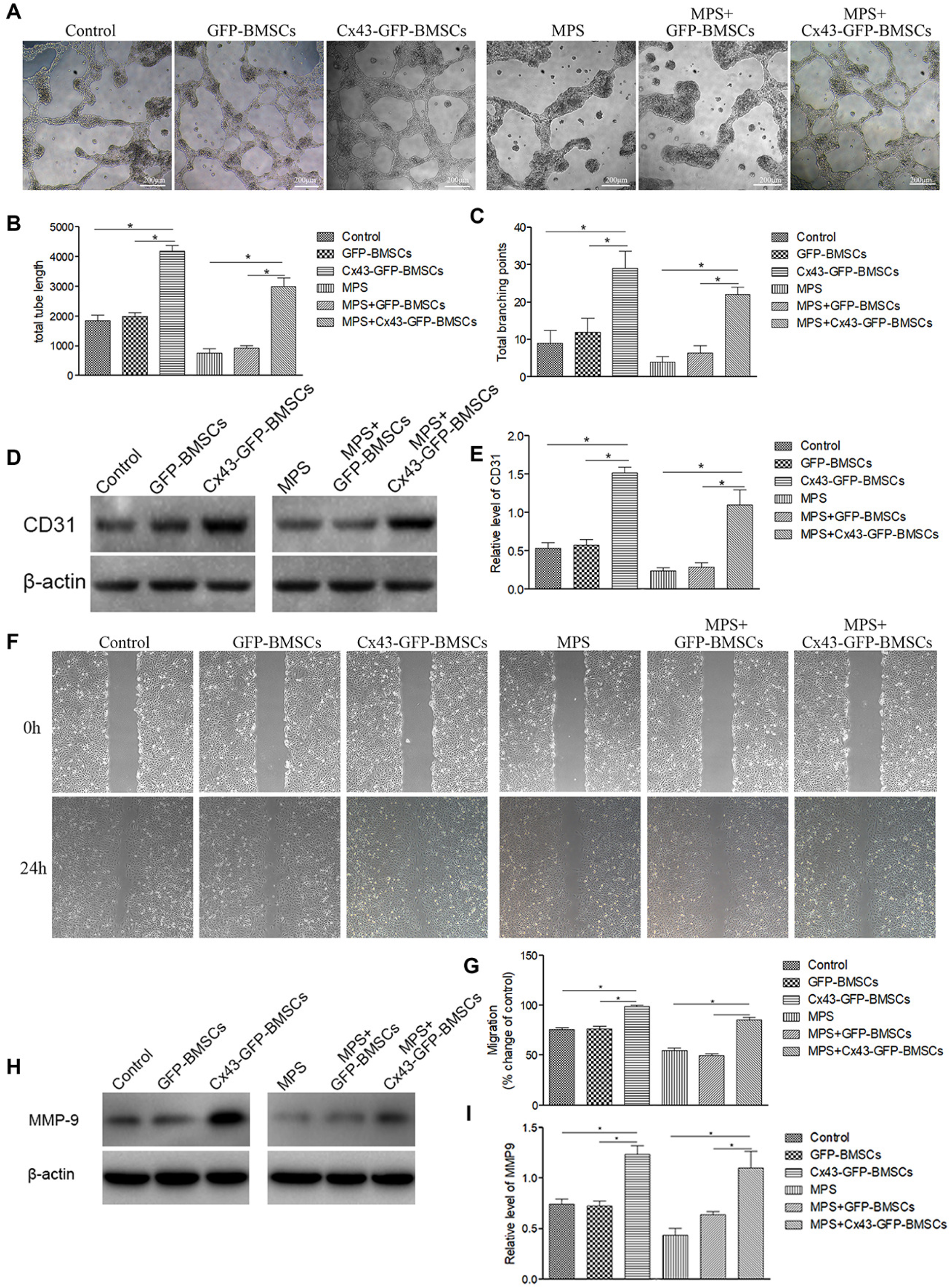
Cx43 overexpression in BMSCs promotes angiogenesis in vitro. (A) Cell supernatants of Cx43-GFP-BMSCs had significantly promoted tube formation compared with GFP-BMSCs or control groups. Tube formation was reduced by MPS, but Cx43 overexpression reversed the effects that were caused by MPS. (B-C) The finding of the changes of total tube length and total branching points were in line with the results that were observed for tube formation. (D) HUVECs were cultured in the supernatants of Cx43-GFP-BMSCs, GFP-BMSCs or non-transfected cells for 4 days, and the expression level for CD31 was detected by Western blot. (E) Statistical analysis of CD31 expression. (F) The supernatants of Cx43-GFP-BMSCs had significantly promoted HUVECs’ migration compared with the GFP-BMSCs or control groups. HUVECs’ migration ability was decreased by MPS treatment, whereas Cx43 overexpression reversed the effects that were caused by MPS; (G) Statistical analysis for all the groups of cells migration. (H) The expression of MMP9 in HUVECs was measured by western blot analysis. (I) Statistical analysis for MMP9 expression; Each experiment was repeatedly performed at least three times. *P<0.05.

Consistent with the results that were obtained in the tube formation assay, the supernatants of Cx43-GFP-BMSCs had significantly increased the migration ability of HUVECs cells, in a comparison with the supernatants of GFP-BMSCs or control groups, which was verified by wound-healing assay. MPS decreased the migration ability of HUVECs, and the supernatants of Cx43-GFP-BMSCs reversed these effects of MPS (Fig. 2F, 2G). In addition, the expression of MMP9, which is one of the markers of migration, was detected, and HUVECs were cultured in the supernatants of Cx43-GFP-BMSCs, GFP-BMSCs or non-transfected cells for 4 days. Results demonstrated that the expression of MMP9 was significantly increased under the treatment of the Cx43-GFP-BMSC supernatants in both the absence or presence conditions of MPS (Fig. 2H, 2I). Therefore, these results indicate that Cx43 overexpression could promote angiogenesis in vitro.

### Osteogenic-associated proteins were up-regulated after Cx43 overexpression in BMSCs

ALP staining was used to evaluate cells osteogenic differentiation, and the results showed that the osteogenic differentiation ability was obviously increased after transfected with Cx43-GFP-BMSCs in a comparison with the GFP-BMSCs and control groups in both the absence or presence conditions of MPS (Fig. 3A). Alizarin red staining indicated that more calcium nodules were observed in the Cx43-GFP-BMSCs group than in the GFP-BMSCs and control groups (Fig. 3B). Fewer calcium nodules were observed after induced by MPS; however, the number of calcium nodules was significantly increased in the group of Cx43 overexpressing in BMSCs under the treatment of MPS(Fig. 3B). Osteogenic related proteins, including Runx2, ALP, Collagen I (COL I) and OCN, play an essential role in the process of promoting osteogenesis. Our results showed that the protein expressions levels of Runx2, ALP, and COL I were significantly up-regulated in the Cx43 overexpression group at protein levels when compared with the GFP-BMSC group and the control group. Additionally, Dex could reduce the expressions of Runx2, ALP, and COL I, while Cx43 overexpression in BMSCs had reversed the inhibition effects of MPS (Fig.3C-3F). Furthermore, the expressions levels of Runx2, ALP, COL I were also detected by immunofluorescence method, and the results were consistent with the western blot results (Fig 4). All the findings above showed that the Cx43 overexpression in BMSCs could facilitate the osteogenic differentiation in vitro.

**Figure 3.**
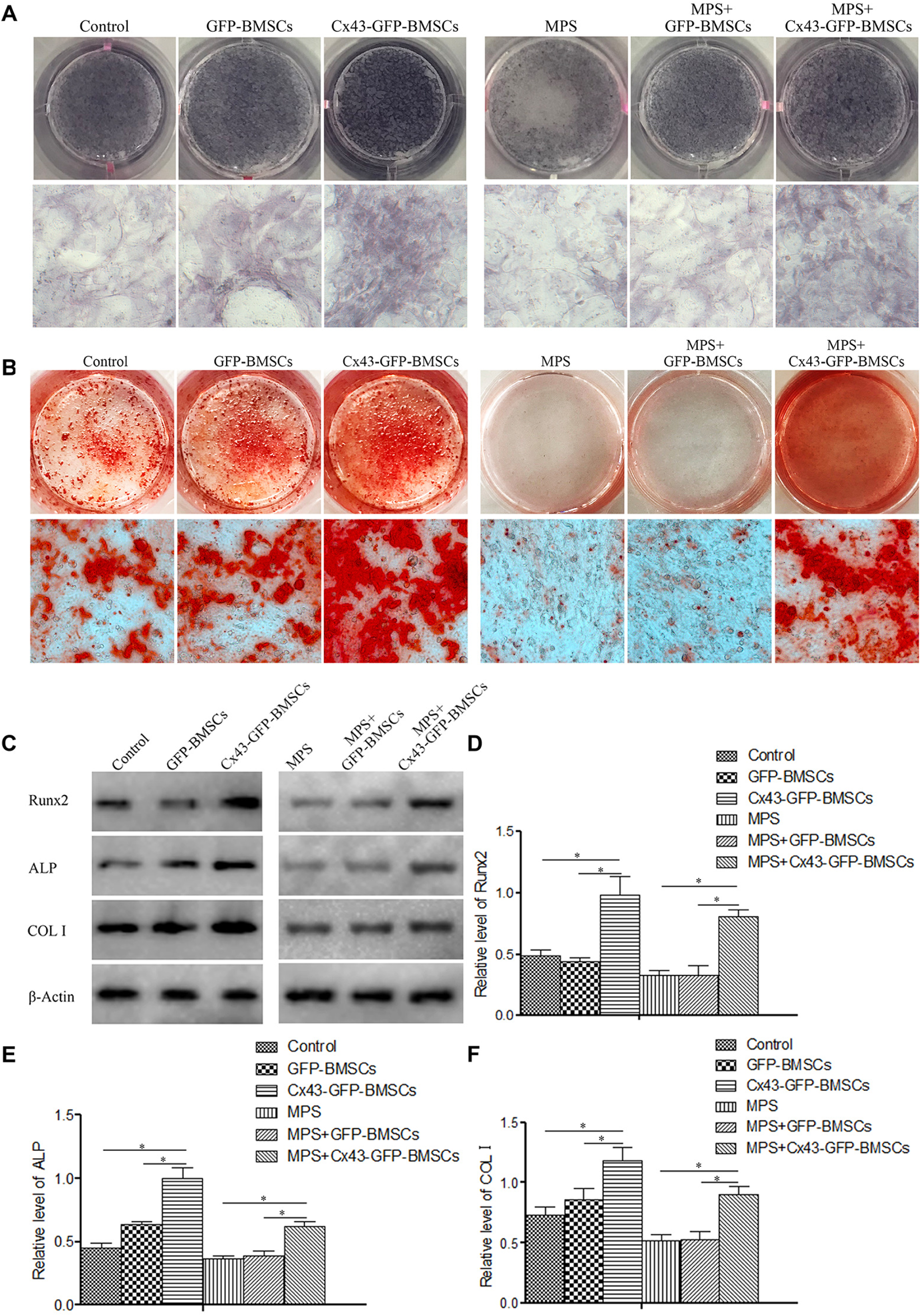
Cx43 overexpression had significantly promoted calcium nodules formation and the expression of osteogenic-related proteins in BMSCs in vitro. (A) The expression of ALP was determined by ALP staining. ALP expression was significantly increased in the Cx43-GFP-BMSCs group, and the ALP expression that was decreased by MPS was reversed by Cx43 overexpression in BMSCs. (B) Cx43-GFP-BMSCs induced the formation of a higher number of calcium nodules than that in the GFP-BMSCs, which was identified by Alizarin red staining. The number of calcium nodules was decreased by MPS and was obviously reversed by Cx43 overexpression in BMSCs. (C) The expressions levels of osteogenic-related proteins, including Runx2, ALP, and COL I were upregulated at the protein level in BMSCs after Cx43 overexpression. MPS decreased the expressions of Runx2, ALP, and COL I, and their expressions were restored by Cx43 overexpression. (D) Statistical analysis for Runx2 expression. (E) Statistical analysis for ALP expression. (F) Statistical analysis for COLI expression; Each experiment was performed repeatedly at least three times. *P<0.05.

**Figure 4.**
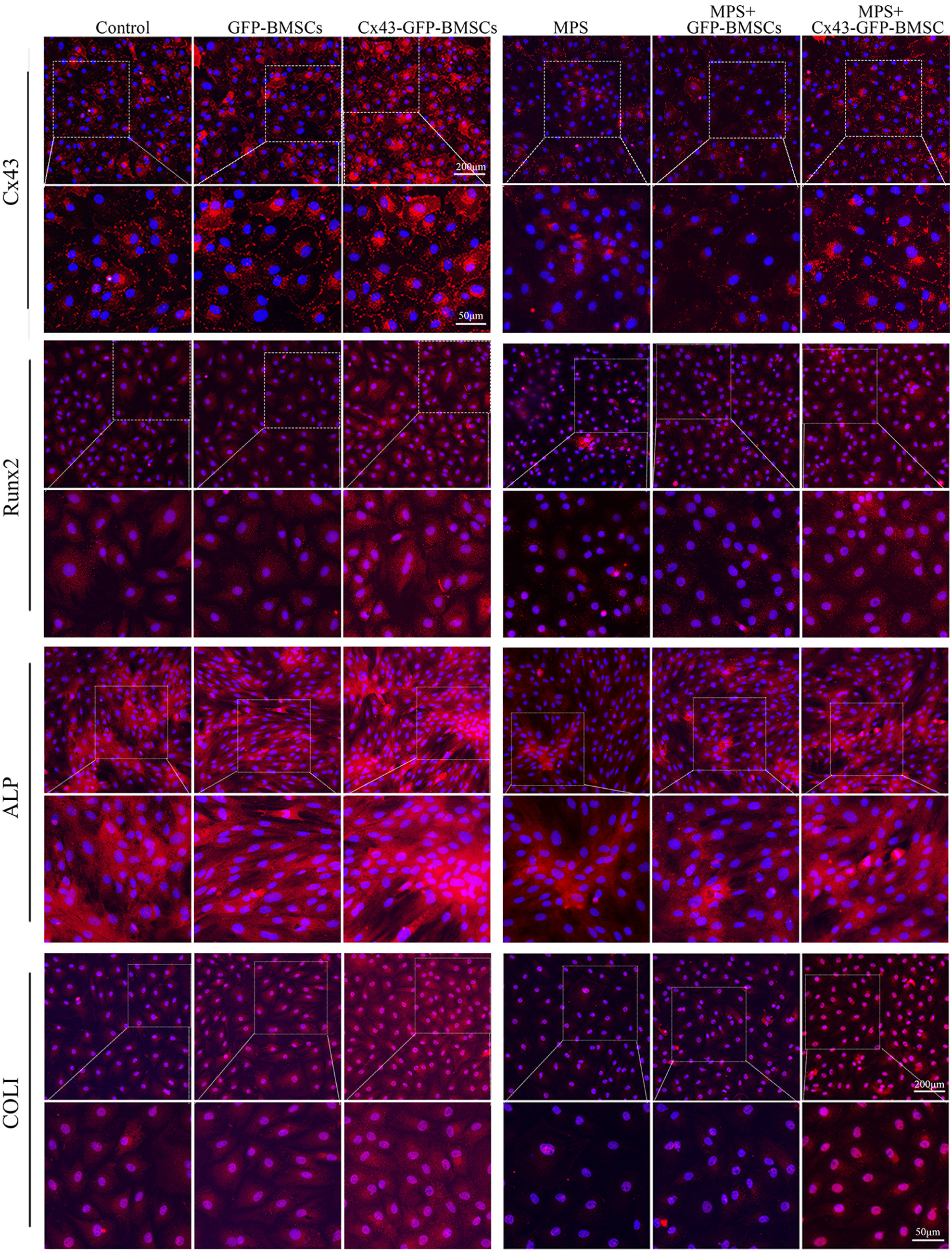
Immunofluorescence staining analysis about the expression of Cx43, Runx2, ALP, and COL I in normal or transfected BMSCs treated or untreated with MPS. (A) Double immunofluorescence staining of Cx43 (red) and DAPI (blue); (B) Double immunofluorescence staining of Runx2 (red) and DAPI (blue); (C) Double immunofluorescence staining of ALP (red) and DAPI (blue); (D) Double immunofluorescence staining of COL I (red) and DAPI (blue). Magnified area in the frame showed that the fluorescence intensity of Cx43, Runx2, ALP and COL I were significantly increased in the Cx43-GFP-BMSCs group, and all of the above indexes of fluorescence intensity were decreased by MPS, which were reversed by Cx43 overexpression in BMSCs. Images are representatives of at least three experiments.

### Cx43 overexpression in BMSCs accelerates osteogenesis in a GIONFH rat model

To further study the role of Cx43 in ONFH in vivo, a rat model of GIONFH was successfully established and was identified by histomorphology as described previously. The results of H&E staining indicated that Cx43 overexpression in BMSCs reduced the morphological changes that were induced by MPS in vivo (Fig. 5A). In addition, our results demonstrated that BMSCs transplantation had partially promoted osteogenesis in GIONFH, whereas Cx43 overexpression of BMSCs had significantly improved osteogenesis (Fig. 5B). The expressions levels of Runx2, ALP, COL I were decreased by MPS treatment at the protein level, and their expressions were rescued by the transplantation with Cx43 overexpression BMSCs in the femoral head tissues (Fig. 5C-5E). The presence of the transplanted BMSCs was confirmed by GFP immunohistochemical staining, and the results showed that the transgenic BMSCs were successfully located in the femoral head (Fig. 6A). The trabecular changes in the subchondral area of the femoral heads were visualized by micro-CT scanning. After eight weeks from the first MPS injection, the occurrence rate of empty lacuna was significantly lower in the group of Cx43 overexpressing BMSCs transplantation than in GFP-BMSC group or control group (Fig. 6B). Furthermore, the BV/TV values were significantly decreased in the MPS group, whereas the Cx43 overexpression of BMSCs transplantation had significantly reduced the effect of MPS (Fig. 6C). In addition, the Tb. Th, and Tb. N, were remarkably improved after Cx43-GFP-BMSCs transplantation compared with the GFP-BMSCs group or MPS alone (Fig. 6D, 6E), however, Tb. Sp was significantly decreased after Cx43-GFP-BMSCs transplantation (Fig. 6F). All the results above indicated that Cx43 overexpression of BMSCs transplantation could decrease the osteonecrosis of GIONFH in vivo.

**Figure 5.**
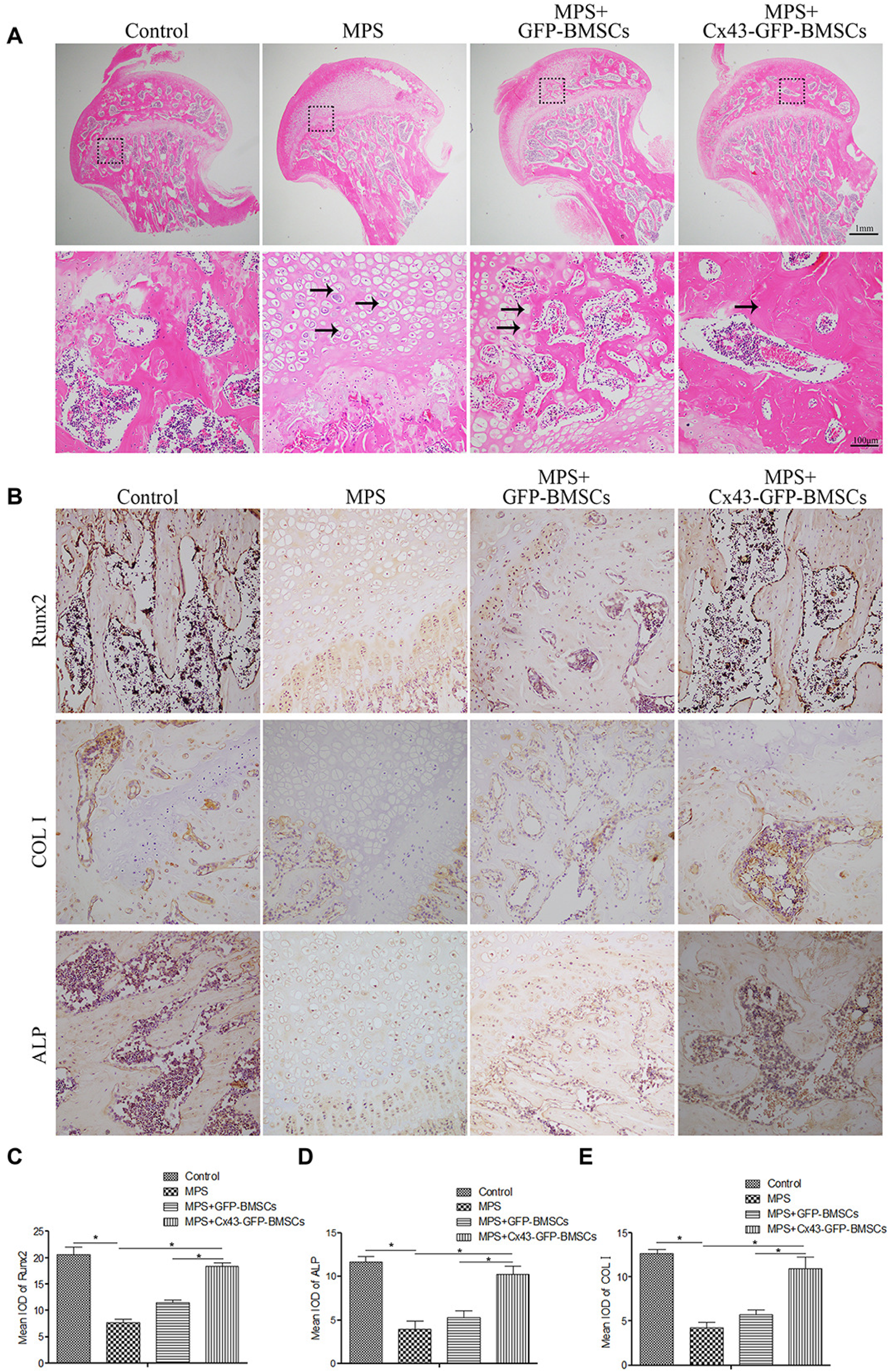
Cx43-GFP-BMSCs promote osteogenesis in vivo. (A) The overexpression of Cx43 in BMSCs reduced the morphological changes that were induced by MPS as indicated by H&E staining. Arrow indicates empty lacunae. (B) Cx43-GFP-BMSCs promoted the expression of Runx2, ALP and COL I despite the presence of MPS. Arrows indicate the expressions of Runx2, ALP and COL I. (C) Density evaluation of Runx2. (D) Density evaluation of ALP. (E) Density evaluation of COL. Each experiment was repeated performed at least three times. *P<0.05.

**Figure 6.**
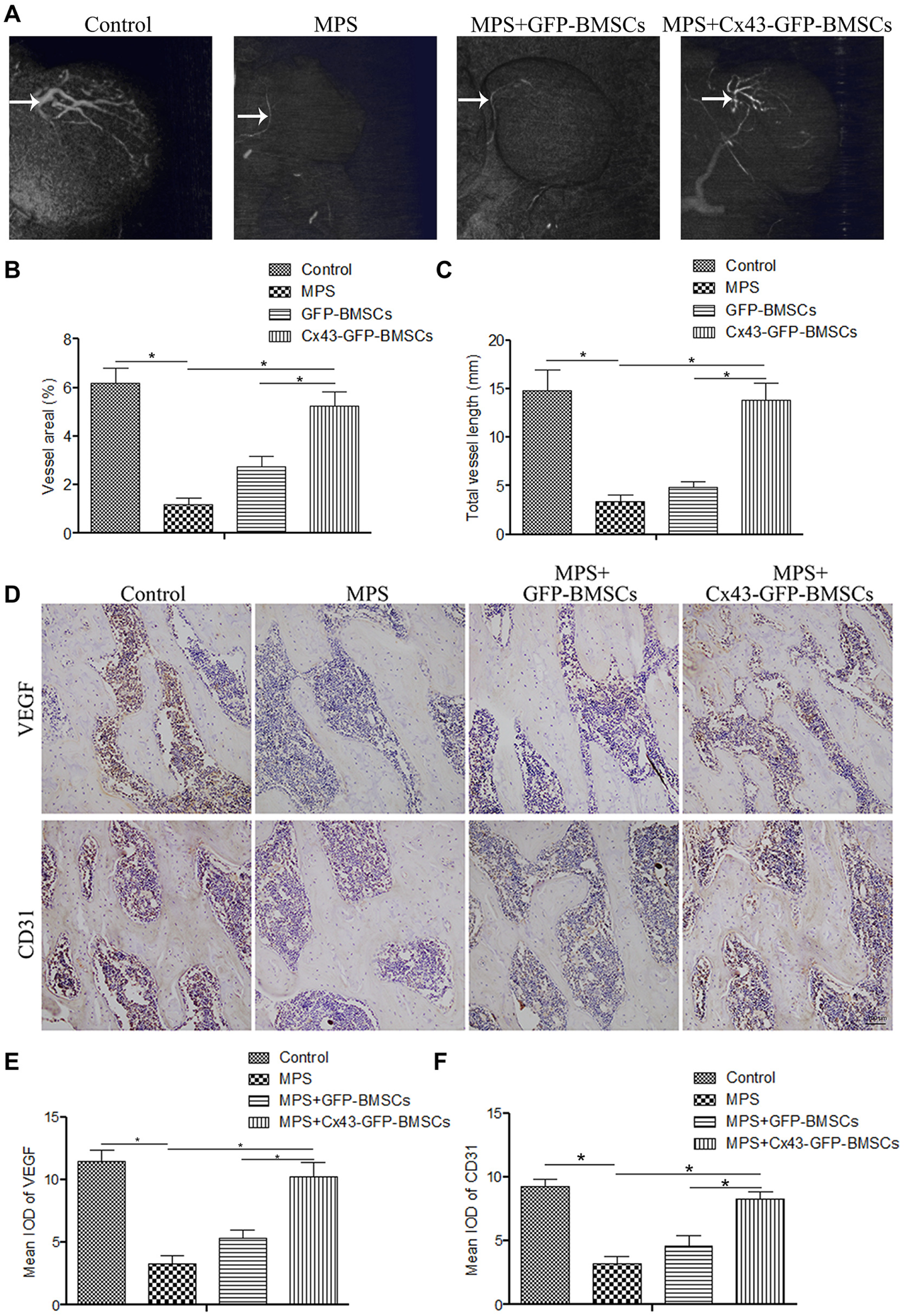
Cx43-GFP-BMSCs promote angiogenesis in vivo. (A) Cx43 overexpression in BMSCs promote vascularization, as indicated by angiography. Arrows indicate vessels in the bones. (B-C) Quantitative analysis of the vascularized area in the femoral head receiving different treatment. (D) The expression of the angiogenesis indicator VEGF and CD31 were upregulated by Cx43-GFP-BMSCs despite the presence of MPS. Arrow indicates the expression of VEGF and CD31; (E) Density evaluation of VEGF; (F) Density evaluation of CD31. Each experiment was repeated performed at least three times. *P<0.05.

### Cx43 overexpression promotes Angiogenesis

Previous results suggested that Cx43 is greatly involved in the process of angiogenesis, and that is essential for osteogenesis. However, its function in the treatment for GIONFH in animal model is still not been investigated. We used angiography to visualize the angiogenesis process in vivo. As shown in Fig. 7A, MPS did not only impair the structure of the femoral head, but also destroyed the vascularization net around it. In the contrary, Cx43-GFP-BMSCs transplantation improved the angiogenesis and had significantly increased the volume of vessel in a comparison with the GFP-BMSCs and control groups. We have also found that the expression of VEGF and CD31 were significantly decreased in the MPS group, while both expressions levels were reversed in the Cx43-GFP-BMSCs transplantation group (Fig. 7B-7D). Therefore, Cx43 overexpression in BMSCs could accelerate angiogenesis in a rat GIONFH model.

**Figure 7.**
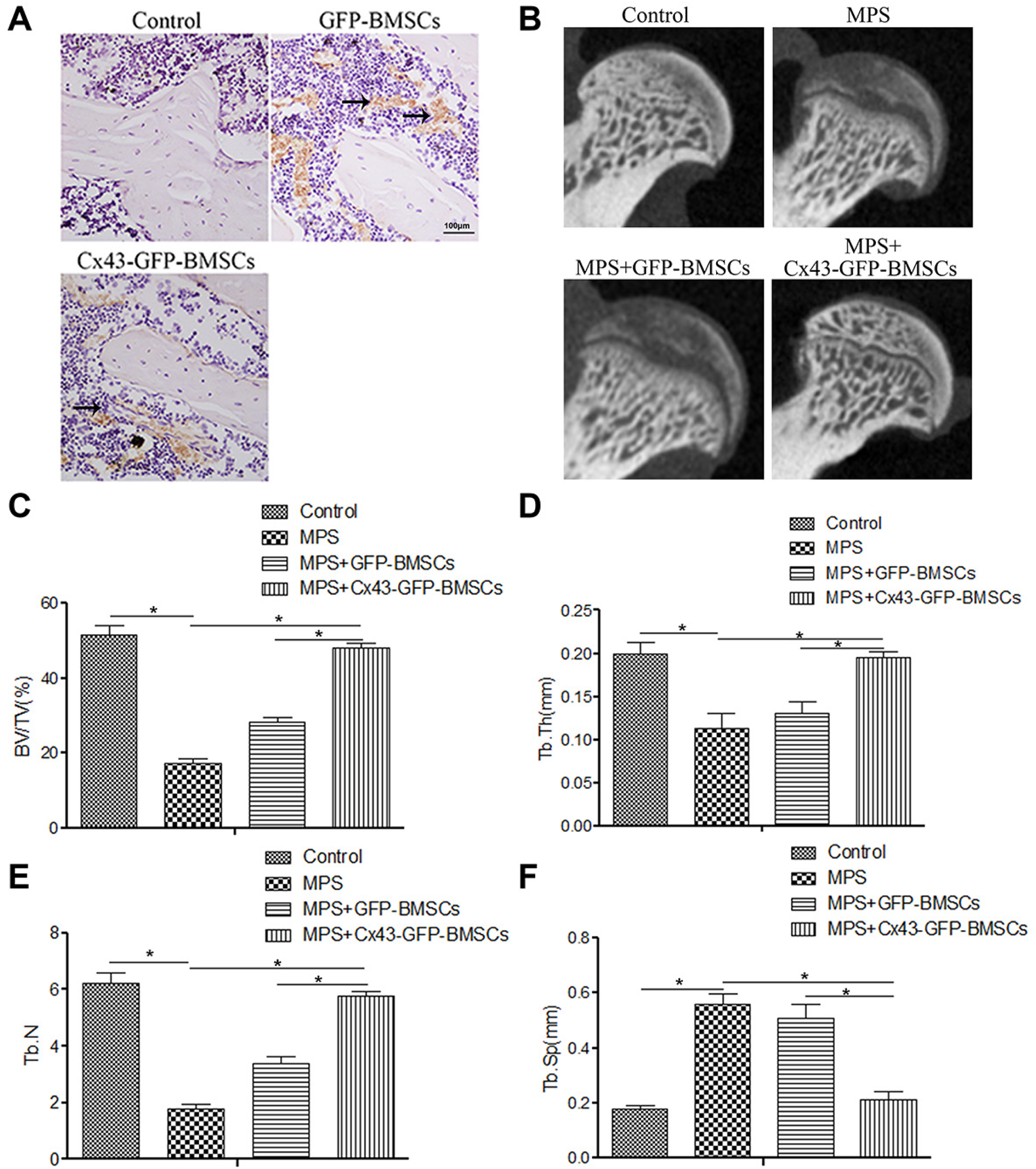
Cx43-GFP-BMSCs promote osteogenesis as indicated by trabecular bone parameters. (A) Immunohistochemistry staining for GFP indicated that the transgenic BMSCs were located in the femoral head. Arrows indicate the localization of GFP-labeled BMSCs. (B) Direct visualization of the morphological changes: Cx43-GFP-BMSCs reversed MPS-induced osteonecrosis. (C-F) Changes of the trabecular bone parameters: bone volume per tissue volume (BV/TV); trabecular thickness (Tb. Th); and trabecular number (Tb. N); trabecular separation (Tb.Sp). Each experiment was repeated performed at least three times. *P<0.05.

## Discussion

Numerous studies have found that there are a lot of factors and molecules that play an important role in BMSC-induced osteogenesis(16, 62), but the definite basic mechanism still has not been explained clearly. We used lentiviral vectors to overexpression Cx43 in BMSCs and found that Cx43 promoted the efficacy of BMSC-induced bone formation by enhancing angiogenesis and osteogenesis. These results showed for the first time that Cx43 might be a vital potential therapeutic target against GIONFH.

Stem cell transplantation has been widely used in the treatment of various diseases, such as alteration of intestinal flora(28), diabetic neuropathy(13), and cardiac disease(34). However, satisfactory therapeutic effects are not always achieved by stem cells transplantation purely. Many studies have suggested that combining stem cells transplantation with genetic modification can improve the treatment efficiency of stem cell transplantation(3, 14, 26). BMSCs transplantation has been proved as a promising treatment method for the early stages of ONFH(10, 38, 52, 58). Previous studies demonstrated that SDF-1α overexpression could significantly promote bone regeneration through osteogenesis and angiogenesis(54), and the overexpression of P-glycoprotein or VEGF165 could also significantly decrease the incidence of GIONFH by promoting osteogenesis and blood vessel regeneration(16–17). In addition, Cx43 was found to be involved greatly in the process of osteogenesis and angiogenesis(35, 49). In this study, we found that Cx43 overexpression in BMSCs had significantly improved the osteogenic differentiation and HUVECs angiogenesis in vitro and promoted bone regeneration by osteogenesis and angiogenesis after cell transplantation in vivo. Our study provided evidence for the first time that Cx43 could promote BMSCs osteogenic differentiation and HUVECs angiogenesis during GIONFH treatment.

Runx2 is an osteogenesis signal molecular that plays an essential role in BMSCs osteogenic differentiation(22, 36). In our work, we found that Runx2 expression was significantly decreased after MPS treatment, whereas the inhibition effects of MPS was attenuated after the overexpression of Cx43 in BMSCs. Lin F et al. reported that the knockdown of Cx43 expression could significantly decrease the osteogenic differentiation of BMSCs, which was verified by the downregulation of Runx2, and this result is in consistent with our findings(32). Osteogenic differentiation of BMSCs is a key physiological process for bone formation, in which the Runx2 signal molecular is involved. As shown in Fig. 3C, we found that Cx43 overexpression could activate Runx2, which suggested that Cx43 could promote osteogenic differentiation in vitro. Furthermore, Micro-CT scanning and H&E staining, directly or indirectly, showed that Cx43 overexpression in BMSCs had significantly reversed MPS-induced osteonecrosis of the femoral head, and that bone trabecular parameters, including BV/TV, Tb. Th and Tb. N were significantly increased after Cx43-GFP-BMSCs transplantation. Therefore, transplantation with Cx43 overexpressing BMSCs could accelerate bone regeneration in GIONFH treatment.

In addition to osteogenesis, BMSCs could also enhance the process of angiogenesis and blood vessel regeneration, which is essential for bone regeneration(37). Recent studies have shown that Cx43 can not only participates in cell migration, but also could promote angiogenesis in many kinds of tissues(11, 41, 46). In this study, we found that Cx43 overexpression in BMSCs could promoted tube formation and HUVECs migration in vitro. In addition, we furtherly confirmed that Cx43-GFP-BMSCs had significantly increased the vascularization and angiogenesis, as well as the volume of vessels, which were verified by angiography. All the results above suggested that Cx43 overexpression in BMSCs could promote BMSCs angiogenesis both in vitro and in vivo, indicating that Cx43 plays an essential role in the process of angiogenesis under the disease of ONFH.

In summary, our results indicate that Cx43 overexpression in BMSCs has good therapeutic effects on GIONFH by promoting angiogenesis and osteogenesis. Additionally, Cx43 is a potential therapeutic molecular for the treatment of GIONFH. However, further studies are still needed to investigate the function of Cx43 in larger animal models of GIONFH.

## Author Contributions

X.Z., and C.C. performed the experiments, analyzed the data, and wrote the manuscript. Y.L. and D.L. carried out the experiments. Q.W. performed the data collection. P.K. and Y.F. study design, critical appraisal of manuscript. All authors approved the final version of the manuscript to be published.

## Author details

^1^Department of Orthopaedics, Shandong Provincial Hospital Affiliated to Shandong First Medical University, Jinan 250021, People’s Republic of China

^2^Department of Orthopaedics, West China Hospital, Sichuan University, No. 37 Wainan Guoxue Road, Chengdu 610041, People’s Republic of China

^3^Pediatric Genetics, Shandong Maternal and Child Health Hospital, Jinan 250021, People’s Republic of China

## Competing Interests

The authors have declared that no competing interest exists.

## Availability of data and materials

The data used to support the findings of this study are available from the corresponding author upon request.

## Consent for publication

All co-authors have read the manuscript and approved its submission to the Molecular and Cellular Biology.

## Ethics approval and consent to participate

All animal experiments were performed based on the guidelines that are formulated by the National Institution of Health on the humane use and care of laboratory animals, and all animal protocols were approved by the Institutes Animal Care and Use Committee of West China Medical School of Sichuan University. This article does not contain any studies with patients who were performed by any of the authors.

## Funding

This work was supported by grants 81974333, 82172414 from the National Natural Science Foundation of China.

## Abbreviations

BMSC: bone-marrow-derived mesenchymal stem cell
Cx43: connexin43
GIONFH: Glucocorticoid induced osteonecrosis of the femoral head
ALP: alkaline phosphatase
COL I: collagen type I
FBS: fetal bovine serum
MPS: methylprednisolone
LPS: lipopolysaccharide
GFP: green fluorescent protein
HUVECs: human umbilical vein endothelial cells
DAPI: 4’6-diamidino-2-phenylindole
SD: standard deviation.

## References

1. Aasen, T., E. Leithe, S.V. Graham, P. Kameritsch, M.D. Mayán, M. Mesnil, K. Pogoda, and A. Tabernero. 2019. Connexins in cancer: bridging the gap to the clinic. Oncogene. 38:4429–4451.

2. Bennett, M.V., J.M. Garré, J.A. Orellana, F.F. Bukauskas, M. Nedergaard, and J.C. Sáez. 2012. Connexin and pannexin hemichannels in inflammatory responses of glia and neurons. Brain Res.. 1487:3–15.

3. Bortolomai, I., M. Sandri, E. Draghici, E. Fontana, E. Campodoni, G.E. Marcovecchio, F. Ferrua, L. Perani, A. Spinelli, T. Canu, M. Catucci, T. Di Tomaso, L. Sergi Sergi, A. Esposito, A. Lombardo, L. Naldini, A. Tampieri, G.A. Hollander, A. Villa, and M. Bosticardo. 2019. Gene Modification and Three-Dimensional Scaffolds as Novel Tools to Allow the Use of Postnatal Thymic Epithelial Cells for Thymus Regeneration Approaches. Stem Cells Transl Med. 8:1107–1122.

4. Cao, Y., J. Xiong, S. Mei, F. Wang, Z. Zhao, S. Wang, and Y. Liu. 2015. Aspirin promotes bone marrow mesenchymal stem cell-based calvarial bone regeneration in mini swine. Stem Cell Res Ther. 6:210.

5. Chuah, Y.J., J.R. Tan, Y. Wu, C.S. Lim, H.T. Hee, Y. Kang, and D.A. Wang. 2020. Scaffold-Free tissue engineering with aligned bone marrow stromal cell sheets to recapitulate the microstructural and biochemical composition of annulus fibrosus. Acta Biomater. 107:129–137.

6. Coutinho, A.E., and K.E. Chapman. 2011. The anti-inflammatory and immunosuppressive effects of glucocorticoids, recent developments and mechanistic insights. Mol. Cell. Endocrinol.. 335:2–13.

7. Czyż, J., K. Piwowarczyk, M. Paw, M. Luty, T. Wróbel, J. Catapano, Z. Madeja, and D. Ryszawy. 2017. Connexin-dependent intercellular stress signaling in tissue homeostasis and tumor development. Acta biochimica Polonica. 64:377–389.

8. Ding, P., W. Zhang, Q. Tan, C. Yao, and S. Lin. 2019. Impairment of circulating endothelial progenitor cells (EPCs) in patients with glucocorticoid-induced avascular necrosis of the femoral head and changes of EPCs after glucocorticoid treatment in vitro. J Orthop Surg Res. 14:226.

9. Ding, Z., and H. Huang. 2015. Mesenchymal stem cells in rabbit meniscus and bone marrow exhibit a similar feature but a heterogeneous multi-differentiation potential: superiority of meniscus as a cell source for meniscus repair. BMC Musculoskelet Disord. 16:65.

10. Döring, M., T. Kluba, K.M. Cabanillas Stanchi, P. Kahle, K. Lenglinger, I. Tsiflikas, C. Treuner, M. Vaegler, M. Mezger, A. Erbacher, M. Schumm, P. Lang, R. Handgretinger, and I. Müller. 2020. Longtime Outcome After Intraosseous Application of Autologous Mesenchymal Stromal Cells in Pediatric Patients and Young Adults with Avascular Necrosis After Steroid or Chemotherapy. Stem Cells Dev...

11. Fan, X., Y. Teng, Z. Ye, Y. Zhou, and W.S. Tan. 2018. The effect of gap junction-mediated transfer of miR-200b on osteogenesis and angiogenesis in a co-culture of MSCs and HUVECs. J. Cell. Sci.. 131.

12. Gangji, V., V. De Maertelaer, and J.P. Hauzeur. 2011. Autologous bone marrow cell implantation in the treatment of non-traumatic osteonecrosis of the femoral head: Five year follow-up of a prospective controlled study. Bone. 49:1005–1009.

13. Gur, S., and W. Hellstrom. 2019. Harnessing Stem Cell Potential for the Treatment of Erectile Function with Diabetes Mellitus: From Preclinical/Clinical Perspectives to Penile Tissue Engineering. Curr Stem Cell Res Ther..

14. Hamed, E.M., M.H. Meabed, U.F. Aly, and R. Hussein. 2019. Recent Progress in Gene Therapy and Other Targeted Therapeutic Approaches for Beta Thalassemia. Curr Drug Targets. 20:1603–1623.

15. Han, L., B. Wang, R. Wang, S. Gong, G. Chen, and W. Xu. 2019. The shift in the balance between osteoblastogenesis and adipogenesis of mesenchymal stem cells mediated by glucocorticoid receptor. Stem Cell Res Ther. 10:377.

16. Han, N., Z. Li, Z. Cai, Z. Yan, Y. Hua, and C. Xu. 2016. P-glycoprotein overexpression in bone marrow-derived multipotent stromal cells decreases the risk of steroid-induced osteonecrosis in the femoral head. J. Cell. Mol. Med.. 20:2173–2182.

17. Hang, D., Q. Wang, C. Guo, Z. Chen, and Z. Yan. 2012. Treatment of osteonecrosis of the femoral head with VEGF165 transgenic bone marrow mesenchymal stem cells in mongrel dogs. Cells Tissues Organs (Print). 195:495–506.

18. Hashimoto, Y., Y. Nishida, S. Takahashi, H. Nakamura, H. Mera, K. Kashiwa, S. Yoshiya, Y. Inagaki, K. Uematsu, Y. Tanaka, S. Asada, M. Akagi, K. Fukuda, Y. Hosokawa, A. Myoui, N. Kamei, M. Ishikawa, N. Adachi, M. Ochi, and S. Wakitani. 2019. Transplantation of autologous bone marrow-derived mesenchymal stem cells under arthroscopic surgery with microfracture versus microfracture alone for articular cartilage lesions in the knee: A multicenter prospective randomized control clinical trial. Regen Ther. 11:106–113.

19. Hernigou, P., A. Dubory, C.H. Flouzat Lachaniette, I. Khaled, N. Chevallier, and H. Rouard. 2018. Stem cell therapy in early post-traumatic talus osteonecrosis. Int Orthop. 42:2949–2956.

20. Hernigou, P., M. Trousselier, F. Roubineau, C. Bouthors, N. Chevallier, H. Rouard, and C.H. Flouzat-Lachaniette. 2016. Stem Cell Therapy for the Treatment of Hip Osteonecrosis: A 30-Year Review of Progress. Clin Orthop Surg. 8:1–8.

21. Hu, Y., S.S. Rao, Z.X. Wang, J. Cao, Y.J. Tan, J. Luo, H.M. Li, W.S. Zhang, C.Y. Chen, and H. Xie. 2018. Exosomes from human umbilical cord blood accelerate cutaneous wound healing through miR-21-3p-mediated promotion of angiogenesis and fibroblast function. Theranostics. 8:169–184.

22. Jiang, H.T., C.C. Ran, Y.P. Liao, J.H. Zhu, H. Wang, R. Deng, M. Nie, B.C. He, and Z.L. Deng. 2019. IGF-1 reverses the osteogenic inhibitory effect of dexamethasone on BMP9-induced osteogenic differentiation in mouse embryonic fibroblasts via PI3K/AKT/COX-2 pathway. J. Steroid Biochem. Mol. Biol.. 191:105363.

23. Jiang, J.X., A.J. Siller-Jackson, and S. Burra. 2007. Roles of gap junctions and hemichannels in bone cell functions and in signal transmission of mechanical stress. Front. Biosci.. 12:1450–1462.

24. Kang, J.S., K.H. Moon, B.S. Kim, D.G. Kwon, S.H. Shin, B.K. Shin, and D.J. Ryu. 2013. Clinical results of auto-iliac cancellous bone grafts combined with implantation of autologous bone marrow cells for osteonecrosis of the femoral head: a minimum 5-year follow-up. Yonsei Med. J.. 54:510–515.

25. Kar, R., N. Batra, M.A. Riquelme, and J.X. Jiang. 2012. Biological role of connexin intercellular channels and hemichannels. Arch. Biochem. Biophys.. 524:2–15.

26. Korshunova, I., S. Rhein, D. García-González, I. Stölting, U. Pfisterer, A. Barta, O. Dmytriyeva, A. Kirkeby, M. Schwaninger, and K. Khodosevich. 2020. Genetic modification increases the survival and the neuroregenerative properties of transplanted neural stem cells. JCI Insight. 5.

27. Kubo, T., K. Ueshima, M. Saito, M. Ishida, Y. Arai, and H. Fujiwara. 2016. Clinical and basic research on steroid-induced osteonecrosis of the femoral head in Japan. J Orthop Sci. 21:407–413.

28. Kusakabe, S., K. Fukushima, T. Maeda, D. Motooka, S. Nakamura, J. Fujita, T. Yokota, H. Shibayama, K. Oritani, and Y. Kanakura. 2020. Pre- and post-serial metagenomic analysis of gut microbiota as a prognostic factor in patients undergoing haematopoietic stem cell transplantation. Br. J. Haematol.. 188:438–449.

29. Leithe, E., M. Mesnil, and T. Aasen. 2018. The connexin 43 C-terminus: A tail of many tales. Biochim Biophys Acta Biomembr. 1860:48–64.

30. Li, J., Q. Shao, X. Zhu, and G. Sun. 2020. Efficacy of autologous bone marrow mesenchymal stem cells in the treatment of knee osteoarthritis and their effects on the expression of serum TNF-α and IL-6. J Musculoskelet Neuronal Interact. 20:128–135.

31. Li, S., H. Zhang, S. Li, Y. Yang, B. Huo, and D. Zhang. 2015. Connexin 43 and ERK regulate tension-induced signal transduction in human periodontal ligament fibroblasts. J. Orthop. Res.. 33:1008–1014.

32. Lin, F.X., G.Z. Zheng, B. Chang, R.C. Chen, Q.H. Zhang, P. Xie, D. Xie, G.Y. Yu, Q.X. Hu, D.Z. Liu, S.X. Du, and X.D. Li. 2018. Connexin 43 Modulates Osteogenic Differentiation of Bone Marrow Stromal Cells Through GSK-3beta/Beta-Catenin Signaling Pathways. Cell. Physiol. Biochem.. 47:161–175.

33. Liu, D., Y. Yang, F. Kuang, S. Qing, B. Hu, and X. Yu. 2019. Risk of infection with different immunosuppressive drugs combined with glucocorticoids for the treatment of idiopathic membranous nephropathy: A pairwise and network meta-analysis. Int. Immunopharmacol.. 70:354–361.

34. Liu, Y., R. Niu, W. Li, J. Lin, C. Stamm, G. Steinhoff, and N. Ma. 2019. Therapeutic potential of menstrual blood-derived endometrial stem cells in cardiac diseases. Cell. Mol. Life Sci.. 76:1681–1695.

35. Moorer, M.C., and J.P. Stains. 2017. Connexin43 and the Intercellular Signaling Network Regulating Skeletal Remodeling. Curr Osteoporos Rep. 15:24–31.

36. Nagel, A.K., and L.E. Ball. 2014. O-GlcNAc modification of the runt-related transcription factor 2 (Runx2) links osteogenesis and nutrient metabolism in bone marrow mesenchymal stem cells. Mol. Cell Proteomics. 13:3381–3395.

37. Nguyen, V.T., B. Canciani, F. Cirillo, L. Anastasia, G.M. Peretti, and L. Mangiavini. 2020. Effect of Chemically Induced Hypoxia on Osteogenic and Angiogenic Differentiation of Bone Marrow Mesenchymal Stem Cells and Human Umbilical Vein Endothelial Cells in Direct Coculture. Cells. 9.

38. Pan, J., Q. Ding, S. Lv, B. Xia, H. Jin, D. Chen, L. Xiao, and P. Tong. 2020. Prognosis after autologous peripheral blood stem cell transplantation for osteonecrosis of the femoral head in the pre-collapse stage: a retrospective cohort study. Stem Cell Res Ther. 11:83.

39. Peng, W.X., and L. Wang. 2017. Adenovirus-Mediated Expression of BMP-2 and BFGF in Bone Marrow Mesenchymal Stem Cells Combined with Demineralized Bone Matrix For Repair of Femoral Head Osteonecrosis in Beagle Dogs. Cell. Physiol. Biochem.. 43:1648–1662.

40. Plotkin, L.I., D.W. Laird, and J. Amedee. 2016. Role of connexins and pannexins during ontogeny, regeneration, and pathologies of bone. BMC Cell Biol.. 17 Suppl 1:19.

41. Pohl, U. 2020. Connexins: Key Players in the Control of Vascular Plasticity and Function. Physiol. Rev. 100:525–572.

42. Rubessa, M., K. Polkoff, M. Bionaz, E. Monaco, D.J. Milner, S.J. Holllister, M.S. Goldwasser, and M.B. Wheeler. 2017. Use of Pig as a Model for Mesenchymal Stem Cell Therapies for Bone Regeneration. Anim. Biotechnol.. 28:275–287.

43. Schepper, J.D., F. Collins, N.D. Rios-Arce, H.J. Kang, L. Schaefer, J.D. Gardinier, R. Raghuvanshi, R.A. Quinn, R. Britton, N. Parameswaran, and L.R. McCabe. 2020. Involvement of the Gut Microbiota and Barrier Function in Glucocorticoid-Induced Osteoporosis. J. Bone Miner. Res.. 35:801–820.

44. Shen, C., M.R. Kim, J.M. Noh, S.J. Kim, S.O. Ka, J.H. Kim, B.H. Park, and J.H. Park. 2016. Glucocorticoid Suppresses Connexin 43 Expression by Inhibiting the Akt/mTOR Signaling Pathway in Osteoblasts. Calcif. Tissue Int.. 99:88–97.

45. Singh, J.A., A. Hossain, A. Kotb, and G. Wells. 2016. Risk of serious infections with immunosuppressive drugs and glucocorticoids for lupus nephritis: a systematic review and network meta-analysis. BMC Med. 14:137.

46. Slavi, N., A.H. Toychiev, S. Kosmidis, J. Ackert, S.A. Bloomfield, H. Wulff, S. Viswanathan, P.D. Lampe, and M. Srinivas. 2018. Suppression of connexin 43 phosphorylation promotes astrocyte survival and vascular regeneration in proliferative retinopathy. Proc. Natl. Acad. Sci. U.S.A.. 115:E5934–5934E5943.

47. Slominski, R.M., R.C. Tuckey, P.R. Manna, A.M. Jetten, A. Postlethwaite, C. Raman, and A.T. Slominski. 2020. Extra-adrenal glucocorticoid biosynthesis: implications for autoimmune and inflammatory disorders. Genes Immun...

48. Tao, S.C., T. Yuan, B.Y. Rui, Z.Z. Zhu, S.C. Guo, and C.Q. Zhang. 2017. Exosomes derived from human platelet-rich plasma prevent apoptosis induced by glucocorticoid-associated endoplasmic reticulum stress in rat osteonecrosis of the femoral head via the Akt/Bad/Bcl-2 signal pathway. Theranostics. 7:733–750.

49. Wang, D.G., F.X. Zhang, M.L. Chen, H.J. Zhu, B. Yang, and K.J. Cao. 2014. Cx43 in mesenchymal stem cells promotes angiogenesis of the infarcted heart independent of gap junctions. Mol Med Rep. 9:1095–1102.

50. Wang, H.H., C.H. Su, Y.J. Wu, J.Y. Li, Y.M. Tseng, Y.C. Lin, C.L. Hsieh, C.H. Tsai, and H.I. Yeh. 2013. Reduction of connexin43 in human endothelial progenitor cells impairs the angiogenic potential. Angiogenesis. 16:553–560.

51. Wang, K., J. Li, Z. Li, B. Wang, Y. Qin, N. Zhang, H. Zhang, X. Su, Y. Wang, and H. Zhu. 2019. Chondrogenic Progenitor Cells Exhibit Superiority Over Mesenchymal Stem Cells and Chondrocytes in Platelet-Rich Plasma Scaffold-Based Cartilage Regeneration. Am J Sports Med. 47:2200–2215.

52. Wu, Z.Y., Q. Sun, M. Liu, B.E. Grottkau, Z.X. He, Q. Zou, and C. Ye. 2020. Correlation between the efficacy of stem cell therapy for osteonecrosis of the femoral head and cell viability. BMC Musculoskelet Disord. 21:55.

53. Xu, S., R. Guo, P.Z. Li, K. Li, Y. Yan, J. Chen, G. Wang, B. Brand-Saberi, X. Yang, and X. Cheng. 2019. Dexamethasone interferes with osteoblasts formation during osteogenesis through altering IGF-1-mediated angiogenesis. J. Cell. Physiol...

54. Yang, F., F. Xue, J. Guan, Z. Zhang, J. Yin, and Q. Kang. 2018. Stromal-Cell-Derived Factor (SDF) 1-Alpha Overexpression Promotes Bone Regeneration by Osteogenesis and Angiogenesis in Osteonecrosis of the Femoral Head. Cell. Physiol. Biochem.. 46:2561–2575.

55. Yu, H., P. Liu, W. Zuo, X. Sun, H. Liu, F. Lu, W. Guo, and Q. Zhang. 2020. Decreased angiogenic and increased apoptotic activities of bone microvascular endothelial cells in patients with glucocorticoid-induced osteonecrosis of the femoral head. BMC Musculoskelet Disord. 21:277.

56. Zha, X., B. Sun, R. Zhang, C. Li, Z. Yan, and J. Chen. 2018. Regulatory effect of microRNA-34a on osteogenesis and angiogenesis in glucocorticoid-induced osteonecrosis of the femoral head. J. Orthop. Res.. 36:417–424.

57. Zhao, D., D. Cui, B. Wang, F. Tian, L. Guo, L. Yang, B. Liu, and X. Yu. 2012. Treatment of early stage osteonecrosis of the femoral head with autologous implantation of bone marrow-derived and cultured mesenchymal stem cells. Bone. 50:325–330.

58. Zhao, D., Y. Liu, C. Ma, G. Gu, and D.F. Han. 2019. A Mini Review: Stem Cell Therapy for Osteonecrosis of the Femoral Head and Pharmacological Aspects. Curr. Pharm. Des.. 25:1099–1104.

59. Zhao, L., S. Jiang, and B.M. Hantash. 2010. Transforming growth factor beta1 induces osteogenic differentiation of murine bone marrow stromal cells. Tissue Eng Part A. 16:725–733.

60. Zhao, X., X.W. Wang, K.S. Zhou, W. Nan, Y.Q. Guo, J.L. Kou, J. Wang, Y.Y. Xia, and H.H. Zhang. 2017. Expression of Ski and its role in astrocyte proliferation and migration. Neuroscience. 362:1–12.

61. Zhao, X., Z. Wei, D. Li, Z. Yang, M. Tian, and P. Kang. 2019. Glucocorticoid Enhanced the Expression of Ski in Osteonecrosis of Femoral Head: The Effect on Adipogenesis of Rabbit BMSCs. Calcif. Tissue Int.. 105:506–517.

62. Zhou, D., Y.X. Chen, J.H. Yin, S.C. Tao, S.C. Guo, Z.Y. Wei, Y. Feng, and C.Q. Zhang. 2018. Valproic acid prevents glucocorticoid-induced osteonecrosis of the femoral head of rats. Int. J. Mol. Med.. 41:3433–3447.

63. Zhun, W., L. Donghai, Y. Zhouyuan, Z. Haiyan, and K. Pengde. 2018. Efficiency of Cell Therapy to GC-Induced ONFH: BMSCs with Dkk-1 Interference Is Not Superior to Unmodified BMSCs. Stem Cells Int. 2018:1340252.

